## Supplementary information for "Distinct stem-like cell populations facilitate functional regeneration of the *Cladonema* medusa tentacle"

### Figure S1

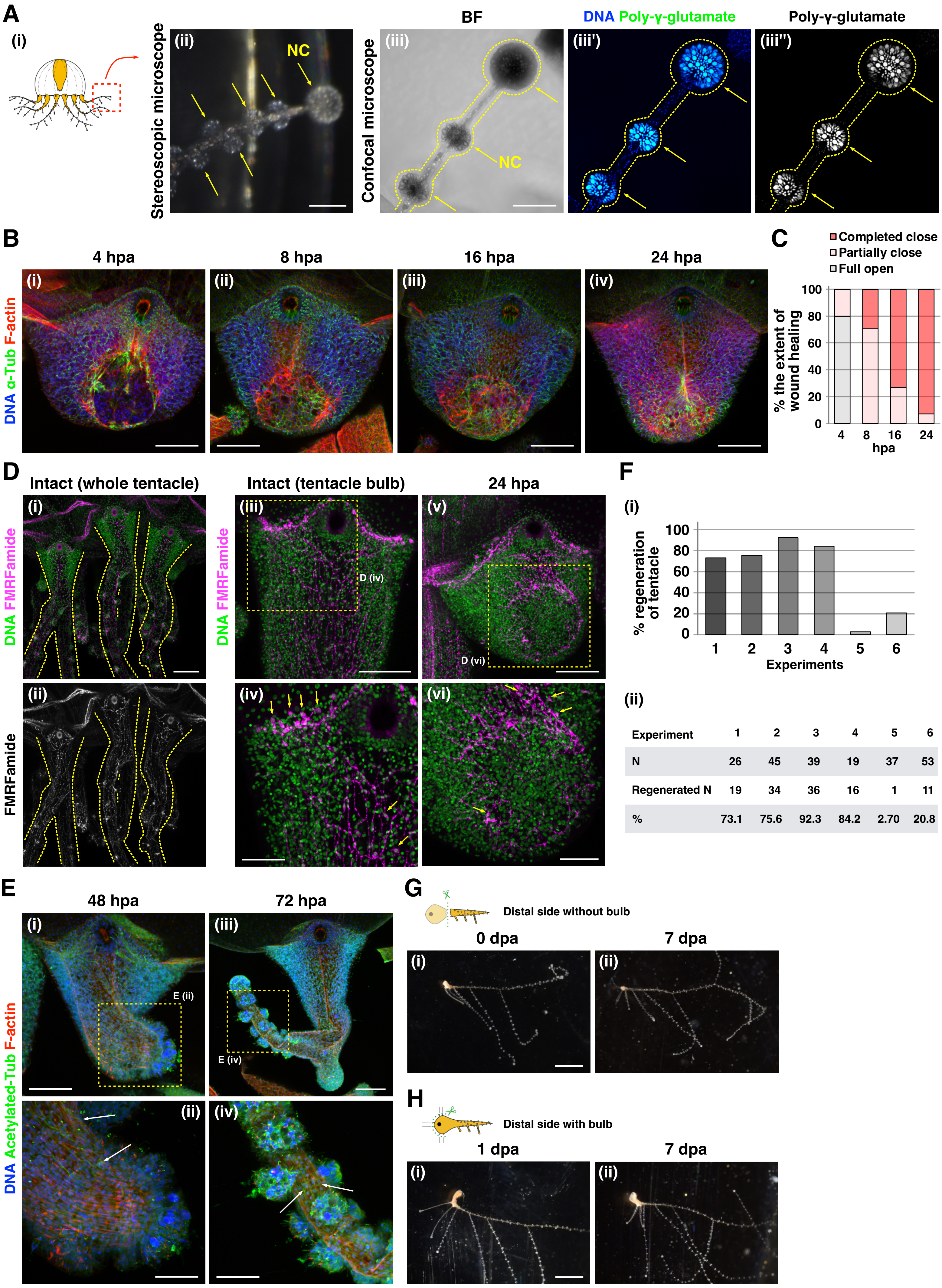


| Experiment | 1 | 2 | 3 | 4 | 5 | 6 |
|---|---|---|---|---|---|---|
| N | 26 | 45 | 39 | 19 | 37 | 53 |
| Regenerated N | 19 | 34 | 36 | 16 | 1 | 11 |
| % | 73.1 | 75.6 | 92.3 | 84.2 | 2.70 | 20.8 |

#### Figure S2

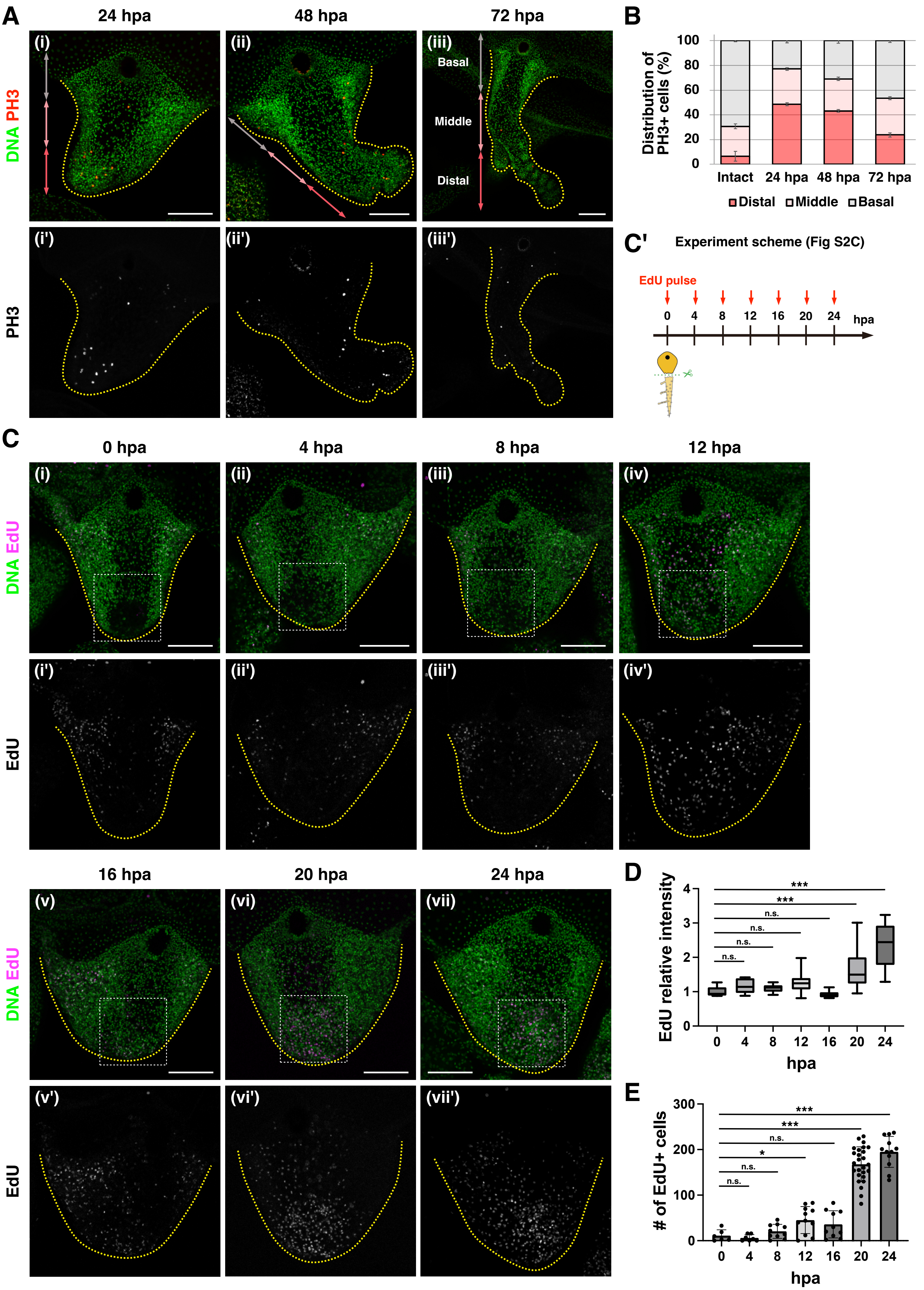

A
24 hpa
48 hpa
72 hpa
DNA PH3
PH3
Basal
Middle
Distal
B
Distribution of PH3+ cells (%)
Intact
24 hpa
48 hpa
72 hpa
Distal
Middle
Basal
C'
Experiment scheme (Fig S2C)
EdU pulse
hpa
C
0 hpa
4 hpa
8 hpa
12 hpa
16 hpa
20 hpa
24 hpa
DNA EdU
EdU
D
EdU relative intensity
hpa
E
### of EdU+ cells
hpa

Figure S3

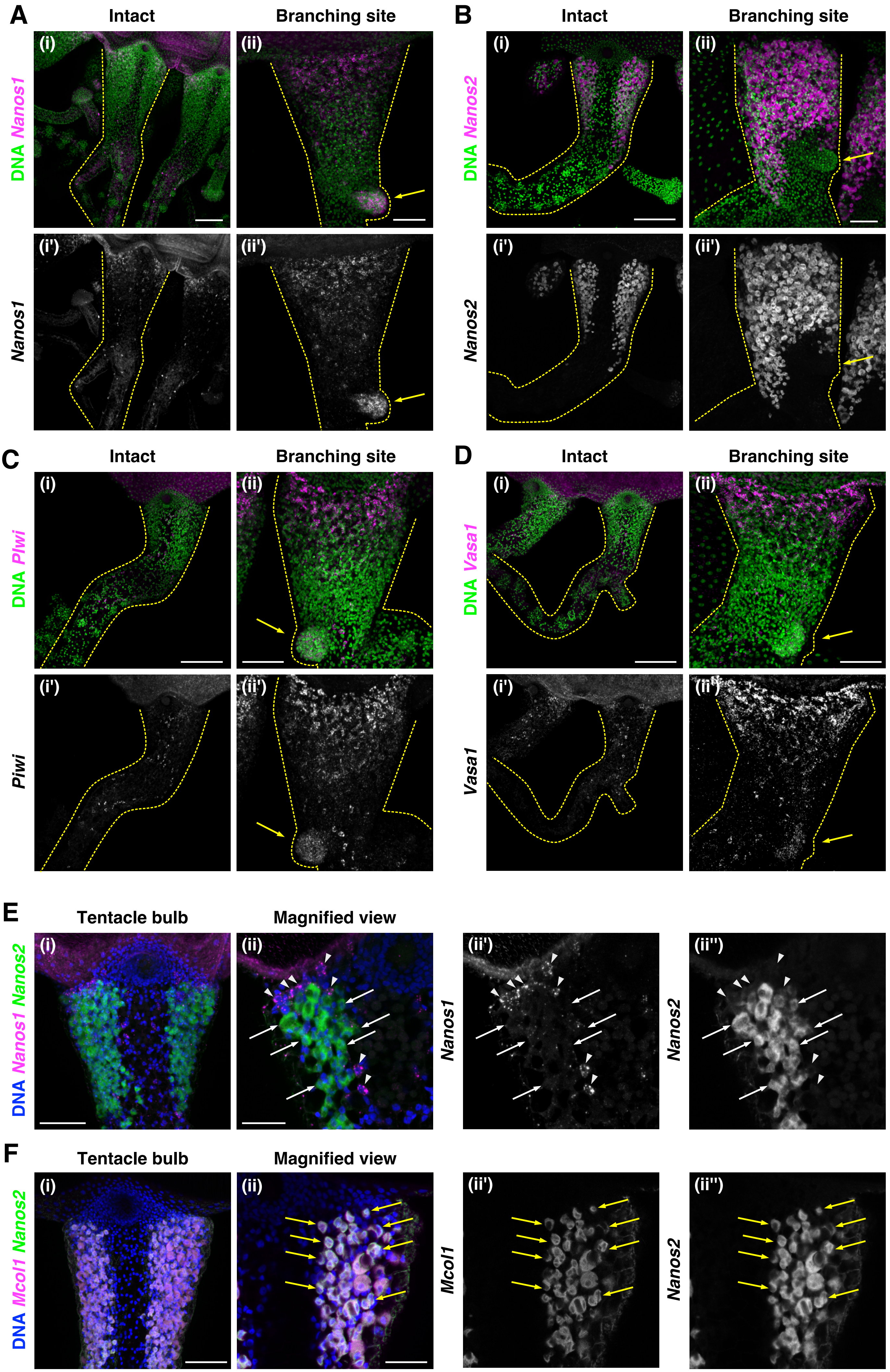
A
Intact
Branching site
(i)
(ii)
(i')
(ii')
DNA Nanos1
Nanos1
B
Intact
Branching site
(i)
(ii)
(i')
(ii')
DNA Nanos2
Nanos2
C
Intact
Branching site
(i)
(ii)
(i')
(ii')
DNA Piwi
Piwi
D
Intact
Branching site
(i)
(ii)
(i')
(ii')
DNA Vasa1
Vasa1
E
Tentacle bulb
Magnified view
(i)
(ii)
(ii')
(ii'')
DNA Nanos1 Nanos2
Nanos1
Nanos2
F
Tentacle bulb
Magnified view
(i)
(ii)
(ii')
(ii'')
DNA Mcol1 Nanos2
Mcol1
Nanos2

### Figure S4

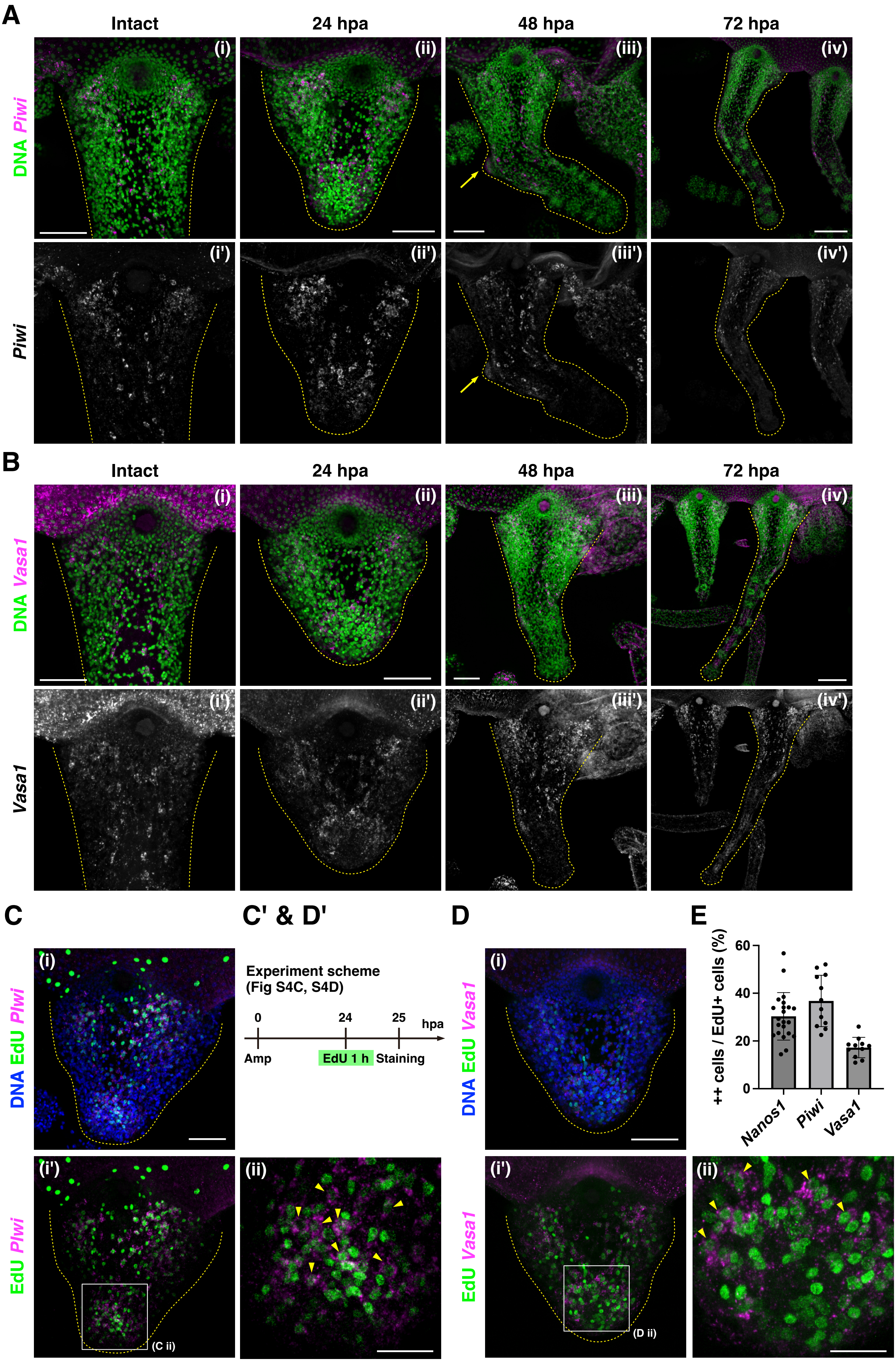

A
Intact
24 hpa
48 hpa
72 hpa
DNA Piwi
Piwi
(i)
(ii)
(iii)
(iv)
(i')
(ii')
(iii')
(iv')
B
Intact
24 hpa
48 hpa
72 hpa
DNA Vasa1
Vasa1
C
DNA EdU Piwi
EdU Piwi
(C ii)
C' & D'
Experiment scheme
(Fig S4C, S4D)
0
24
25
hpa
Amp
EdU 1 h
Staining
D
DNA EdU Vasa1
EdU Vasa1
(D ii)
E
++ cells / EdU+ cells (%)
60
40
20
0
Nanos1
Piwi
Vasa1

### Figure S5

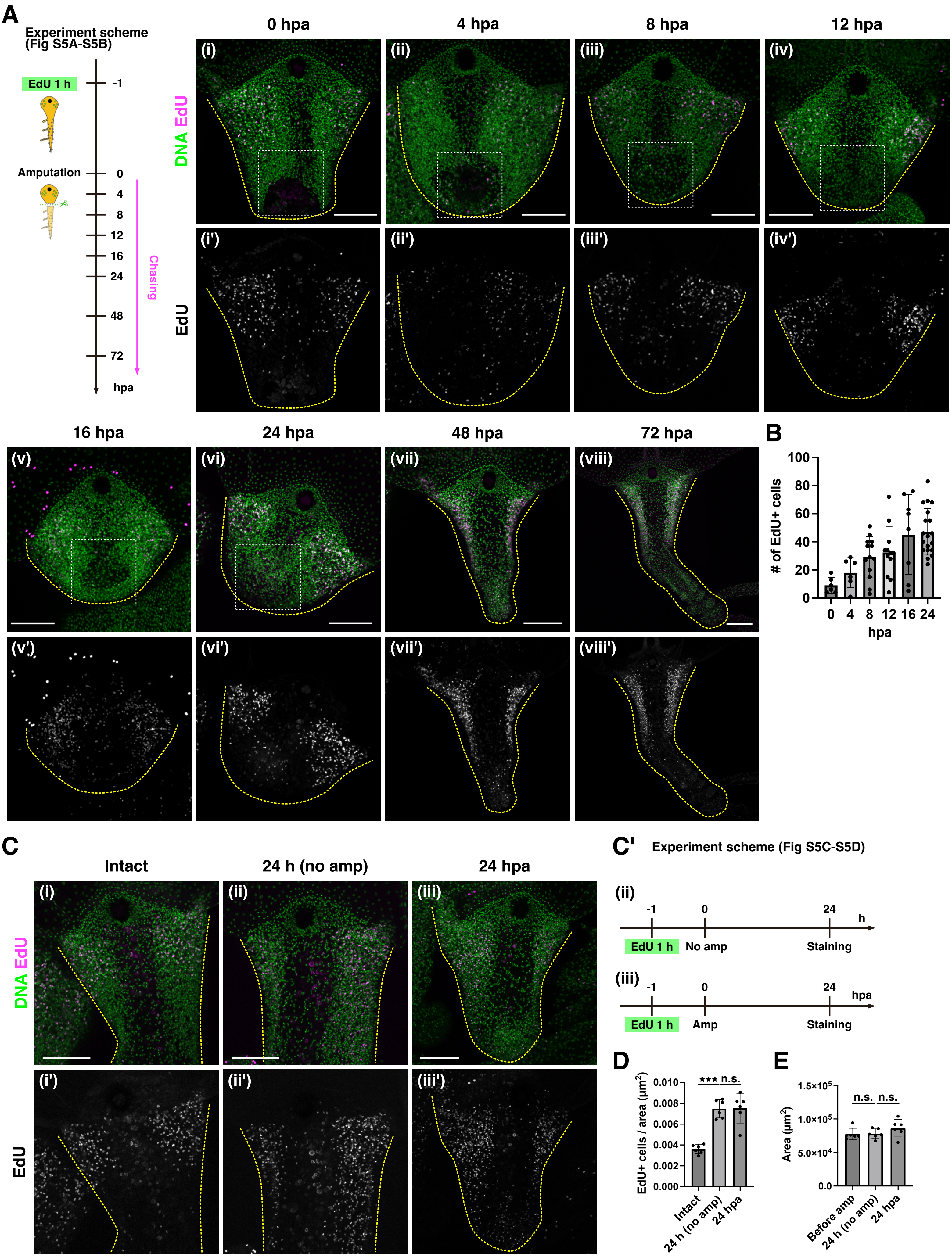
A
Experiment scheme (Fig S5A-S5B)
EdU 1 h
-1
Amputation
0
4
8
12
16
24
48
72
hpa
Chasing
0 hpa
4 hpa
8 hpa
12 hpa
DNA EdU
EdU
(i)
(ii)
(iii)
(iv)
(i')
(ii')
(iii')
(iv')
16 hpa
24 hpa
48 hpa
72 hpa
(v)
(vi)
(vii)
(viii)
(v')
(vi')
(vii')
(viii')
B
### of EdU+ cells
100
80
60
40
20
0
0 4 8 12 16 24
hpa
C
Intact
24 h (no amp)
24 hpa
DNA EdU
EdU
(i)
(ii)
(iii)
(i')
(ii')
(iii')
C'
Experiment scheme (Fig S5C-S5D)
(ii)
-1
0
24
h
EdU 1 h
No amp
Staining
(iii)
-1
0
24
hpa
EdU 1 h
Amp
Staining
D
EdU+ cells / area (μm²)
0.010
0.008
0.006
0.004
0.002
0.000
***
n.s.
Intact
24 h (no amp)
24 hpa
E
Area (μm²)
1.5×10⁵
1.0×10⁵
5.0×10⁴
0.0
n.s.
n.s.
Before amp
24 h (no amp)
24 hpa

### Figure S6

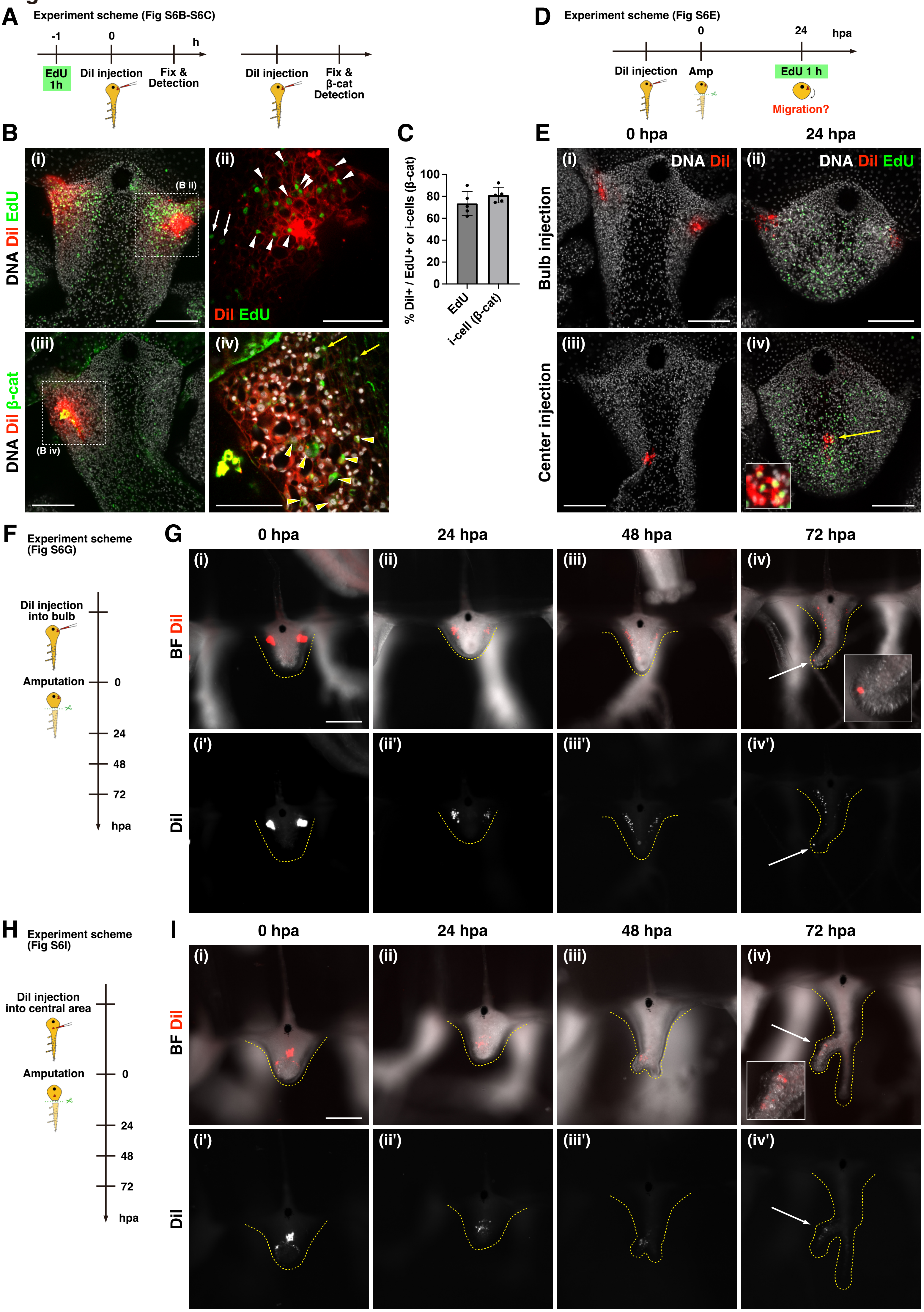
A
Experiment scheme (Fig S6B-S6C)
-1
0
h
EdU 1h
DiI injection
Fix & Detection
DiI injection
Fix & β-cat Detection
D
Experiment scheme (Fig S6E)
0
24
hpa
DiI injection
Amp
EdU 1 h
Migration?
B
DNA DiI EdU
(i)
(B ii)
(ii)
DiI EdU
DNA DiI β-cat
(iii)
(B iv)
(iv)
C
% DiI+ / EdU+ or i-cells (β-cat)
100
80
60
40
20
0
EdU
i-cell (β-cat)
E
0 hpa
24 hpa
Bulb injection
Center injection
(i)
DNA DiI
(ii)
DNA DiI EdU
(iii)
(iv)
F
Experiment scheme (Fig S6G)
DiI injection into bulb
Amputation
0
24
48
72
hpa
G
0 hpa
24 hpa
48 hpa
72 hpa
BF DiI
DiI
(i)
(ii)
(iii)
(iv)
(i')
(ii')
(iii')
(iv')
H
Experiment scheme (Fig S6I)
DiI injection into central area
Amputation
0
24
48
72
hpa
I
0 hpa
24 hpa
48 hpa
72 hpa
BF DiI
DiI
(i)
(ii)
(iii)
(iv)
(i')
(ii')
(iii')
(iv')

### Figure S7

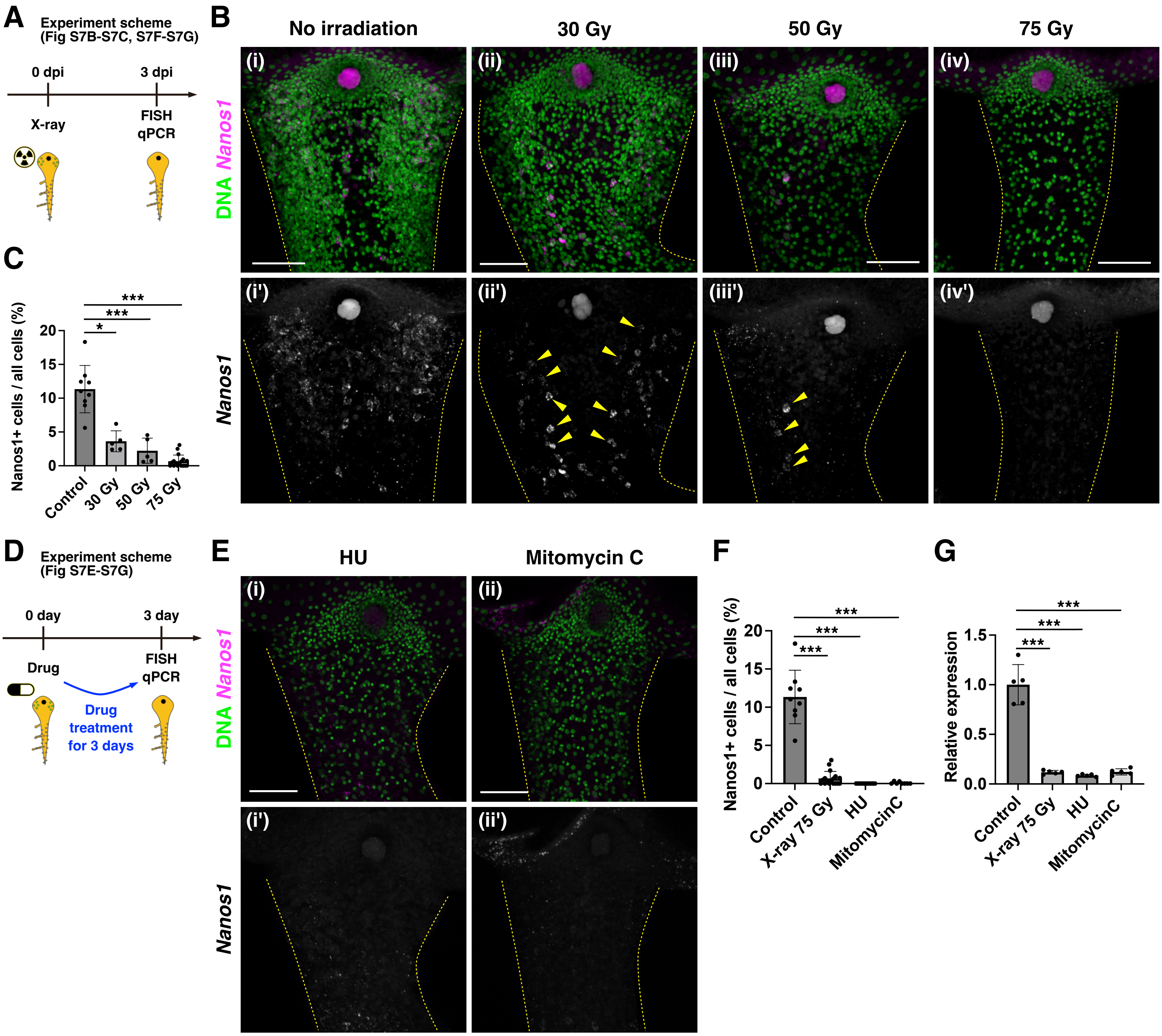
A
Experiment scheme
(Fig S7B-S7C, S7F-S7G)
0 dpi
3 dpi
X-ray
FISH
qPCR
B
No irradiation
30 Gy
50 Gy
75 Gy
(i)
(ii)
(iii)
(iv)
(i')
(ii')
(iii')
(iv')
DNA Nanos1
Nanos1
C
Nanos1+ cells / all cells (%)
0
5
10
15
20
*
***
***
Control
30 Gy
50 Gy
75 Gy
D
Experiment scheme
(Fig S7E-S7G)
0 day
3 day
Drug
FISH
qPCR
Drug
treatment
for 3 days
E
HU
Mitomycin C
(i)
(ii)
(i')
(ii')
DNA Nanos1
Nanos1
F
Nanos1+ cells / all cells (%)
0
5
10
15
20
***
***
***
Control
X-ray 75 Gy
HU
MitomycinC
G
Relative expression
0.0
0.5
1.0
1.5
***
***
***
Control
X-ray 75 Gy
HU
MitomycinC

### Figure S8

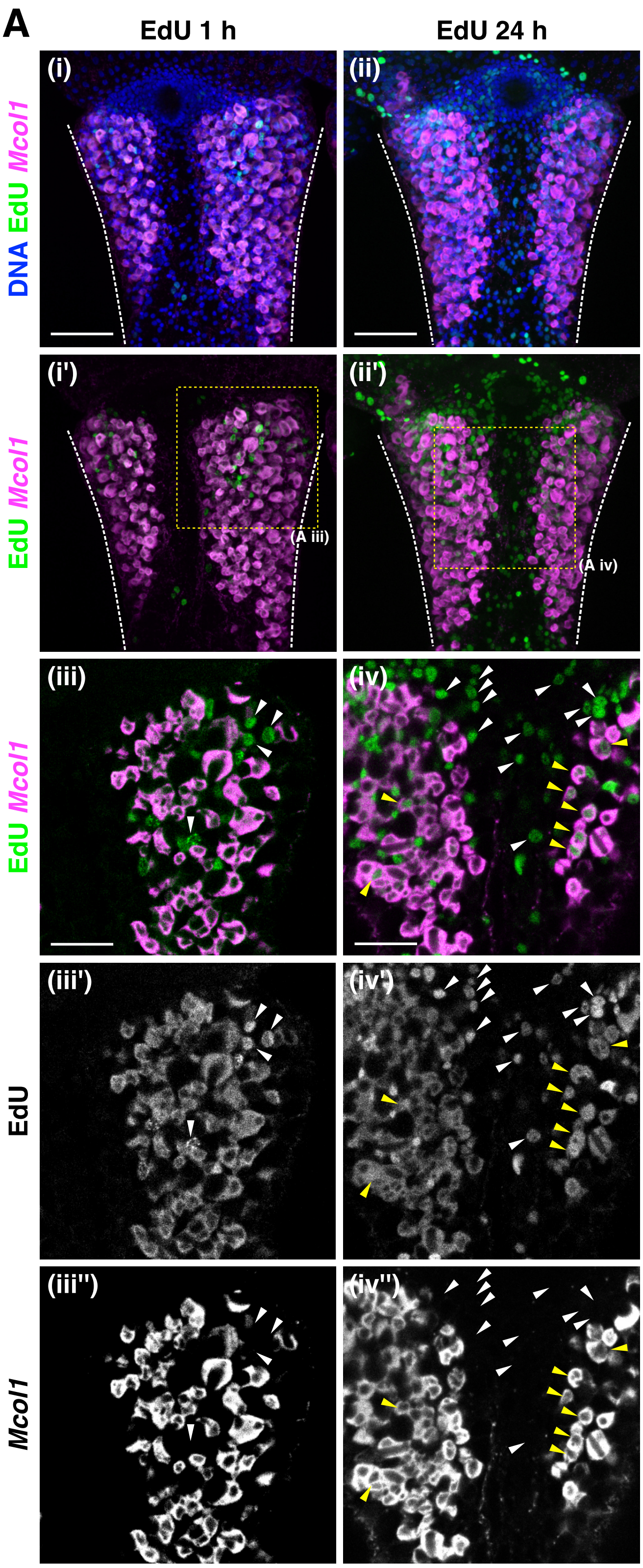
A
EdU 1 h
EdU 24 h
(i)
(ii)
DNA EdU Mcol1
(i')
(ii')
(A iii)
(A iv)
EdU Mcol1
(iii)
(iv)
EdU Mcol1
(iii')
(iv')
EdU
(iii'')
(iv'')
Mcol1

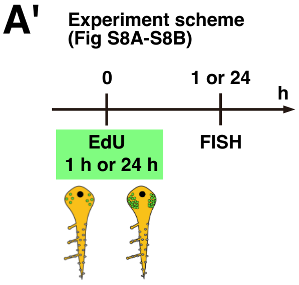
A'
Experiment scheme
(Fig S8A-S8B)
0
1 or 24
h
EdU
1 h or 24 h
FISH

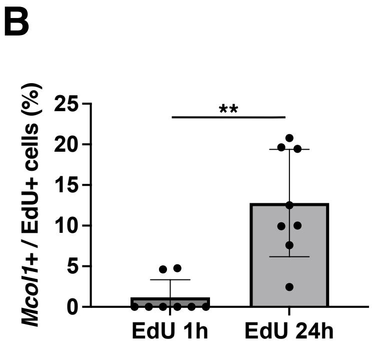
B
Mcol1+ / EdU+ cells (%)
25
20
15
10
5
0
**
EdU 1h
EdU 24h

### Figure S9

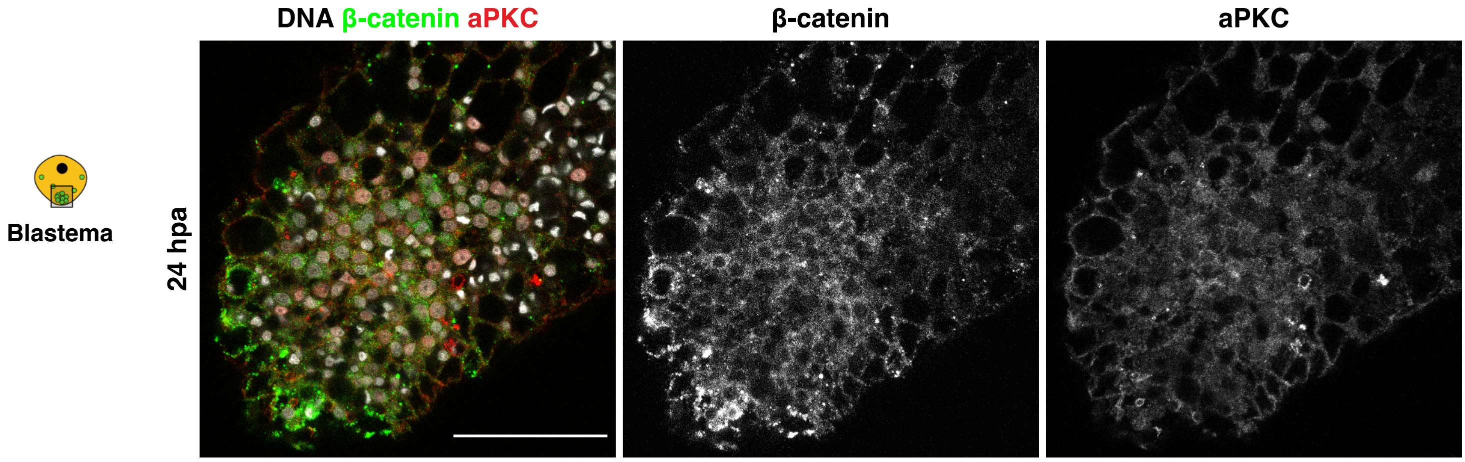

#### Figure S10

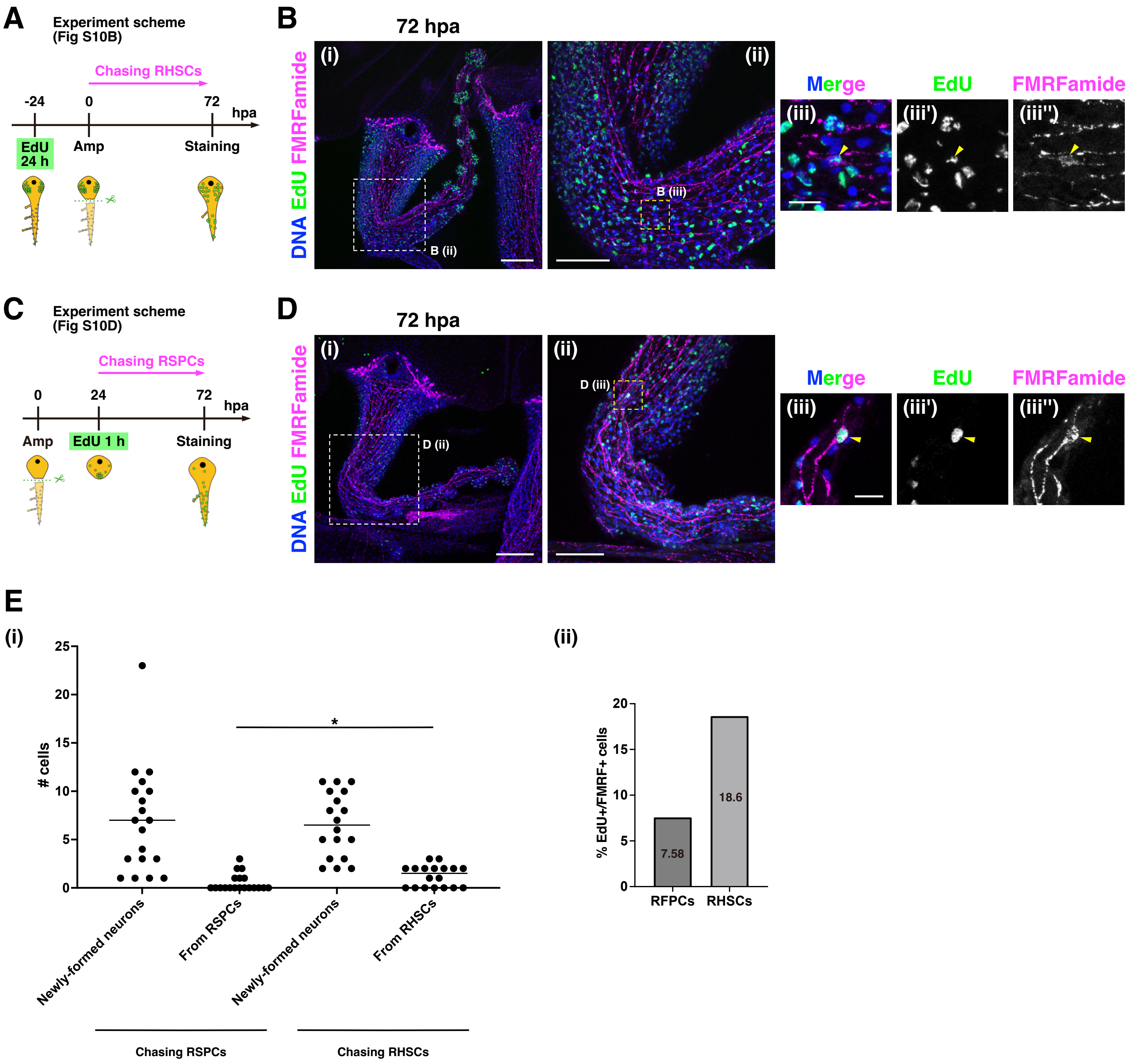
A
Experiment scheme
(Fig S10B)
Chasing RHSCs
-24
0
72
hpa
EdU 24 h
Amp
Staining
B
72 hpa
DNA EdU FMRFamide
(i)
B (ii)
(ii)
B (iii)
Merge
EdU
FMRFamide
(iii)
(iii')
(iii'')
C
Experiment scheme
(Fig S10D)
Chasing RSPCs
0
24
72
hpa
Amp
EdU 1 h
Staining
D
72 hpa
DNA EdU FMRFamide
(i)
D (ii)
(ii)
D (iii)
Merge
EdU
FMRFamide
(iii)
(iii')
(iii'')
E
(i)
### cells
25
20
15
10
5
0
*
Newly-formed neurons
From RSPCs
Newly-formed neurons
From RHSCs
Chasing RSPCs
Chasing RHSCs
(ii)
% EdU+/FMRF+ cells
20
15
10
5
0
7.58
18.6
RFPCs
RHSCs

### Figure S11

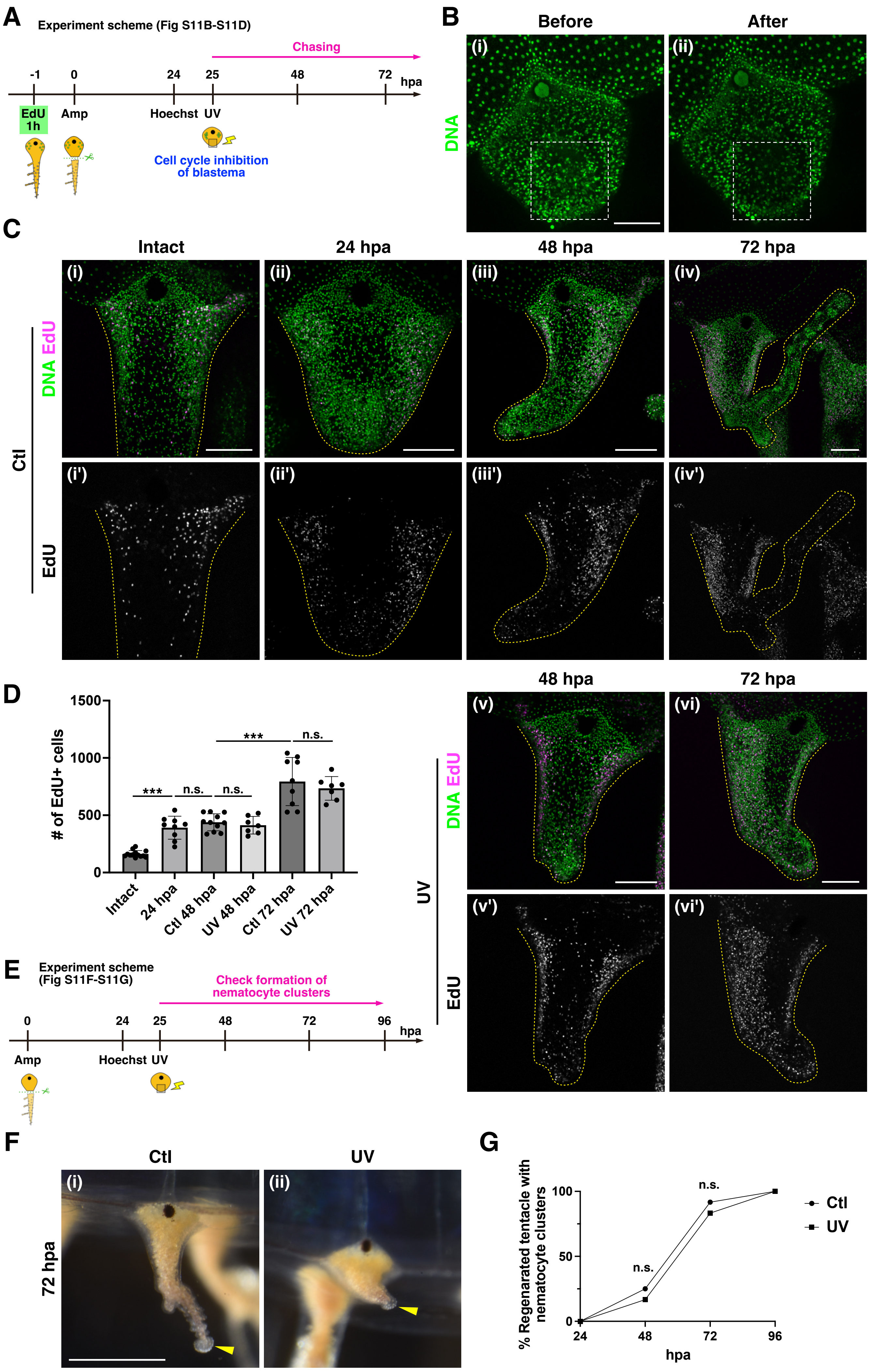
A
Experiment scheme (Fig S11B-S11D)
Chasing
-1
0
24
25
48
72
hpa
EdU 1h
Amp
Hoechst
UV
Cell cycle inhibition of blastema
B
Before
After
(i)
(ii)
DNA
C
Intact
24 hpa
48 hpa
72 hpa
(i)
(ii)
(iii)
(iv)
DNA EdU
Ctl
(i')
(ii')
(iii')
(iv')
EdU
D
1500
1000
500
0
### of EdU+ cells
***
n.s.
n.s.
***
n.s.
Intact
24 hpa
Ctl 48 hpa
UV 48 hpa
Ctl 72 hpa
UV 72 hpa
48 hpa
72 hpa
(v)
(vi)
DNA EdU
UV
(v')
(vi')
EdU
E
Experiment scheme (Fig S11F-S11G)
Check formation of nematocyte clusters
0
24
25
48
72
96
hpa
Amp
Hoechst
UV
F
Ctl
UV
(i)
(ii)
72 hpa
G
% Regenarated tentacle with nematocyte clusters
100
50
0
n.s.
n.s.
24
48
72
96
hpa
Ctl
UV
